# Plants with purple abaxial leaves: A repository of metrics from stomata distribution

**DOI:** 10.1101/294553

**Authors:** Humberto Antunes de Almeida filho, Odemir Martinez Bruno

## Abstract

Plants with purple abaxial leaves are very common in nature but the ecophysiological aspects of this phenotype are not well known. We observed here that the purple color of the abaxial tegument in extant plants make stomata completely visible. Based in it we measured the relations between stomatic density and distance between stomata pairs and observed a general log-normal trend line between density and interstomatic distance. These data shows that measures of stomatic distances at purple abaxial leaves are able to be a sensor to environmental changes of living purple plants. In future ecophysiological inferences will be established from the information brought by the measurements of the distance between stomata in purple plants.

## Introduction

The purple color of lower abaxial leaf surfaces is commonly observed in deeply-shaded understorey plants, especially in the tropical regions. However the physiological role to pigmentation remains unclear [9, 11].The pigments responsible for the coloring are known by anthocyanins, it can be found in plant species across a broad range of habitats, especially common in understorey plants of the tropics [6, 13]. The color of the abaxial surface may be transient or permanent depending of the environmental conditions and plant specie. Although the distribution of the color from abaxial leaves among tropical taxa are widespread, it is not known the physiological roles of this phenotype. Some studies regarding the ecophysiological relationships involved in the foliar anthocyanin in upper adaxial surface have demonstrated that pigmentation play a rule in light-attenuation, protecting underlying cells from photo-inhibition through the absorption of high energy blue-green wavelengths [3]. Photoprotection have also been implicated in the physiological roles played by anthocyanins in plants in which exposed abaxial leaf surfaces are vulnerable to high-incident light [10]. Some studies have observed that the purple abaxial leaf surfaces appeared to reflect more red light than green [14]. This observation induces the hypothesis that anthocyanin pigments may function to reflect adaxially-transmitted red light back into the mesophyll, in order to capture red photons in environments where light is limiting. This hypothesis is known as back-scatter propagation, it has not yet received a complete experimental validation but could explain the color of leaves in understorey plants. It is known that the abaxial surface of leaves in some species of purple plants presents a high concentration of green stomatal cells which generates an interesting contrast between the color of the floor of the epidermis (purple) and the stomata (green), making this cells completely visible in studies of density and distribution pattern on the leaf epidermis of the extant plants [5]. The stomatic density varies according to the environmental light exposure [7], then the color of the epidermis may also influence the distribution of stomata. The Stomatic morphology, distribution and celular physiology respond to a large spectrum of environmental signals and at the same time can be responsible to own climate change [8]. We observed in this work that in plants with purple abaxial leaves, the stomata are completely visible on the extant leaves due to the contrast between the color of the green stomata and the color of the floor, which is purple. This made it possible to propose strategies for measuring the distribution of stomata in the abaxial epidermis. Our approach was based on measuring the distance between pairs of stomata and compiling these measurements into a quadratic matrix of distances. The relation between interstomatic distance and stomatic density showed a general relationship as observed for stomatal density in function of guard cell length as evidenced in previos works [2, 8]. These works demonstrated that the log-normal relationship fitted with mathematical simulations of stomata gaseous exchange models, indicating that the average relationship has virtually the same trend to one in which changes in density and size are exactly compensatory, in terms of *CO*_2_ and water exchange [2, 8]. However, the logarithmic decay between stomatic density as a function of the distance between stomata that we observed in this work has not been demonstrated, especially in purple plants.

Our results show that the phenotypic plasticity of stomatal density can be probed by measurements of the distance between stomata in purple plants. Therefore, ecophysiological inferences regarding environmental changes can also be probed from the phenotypic plasticity of stomatal distribution in plants with purple abaxial leaves, which makes these plants excellent sensors of environmental change.

## 1 Materials and Methods

### 1.1 Optic microscopy

Sections of extant leaves of the plants *Tradescantia pallida, Tradescantia zebrina, Callisia repens, Tradescantia minima, Ctenanthe oppenheimiana* and *Oxalis atropurpurea* were mounted on slides, covered with fresh water and covered with coverslips. Sections of 0.5 *cm*^2^ from the medial third of leaves of the plants *Tradescantia pallida, Tradescantia zebrina* and *Ctenanthe oppenheimiana* were hydrated and subjected to optical microscopy after a maximum period of 60 minutes after excision. Whole leaves of the plants *Callisia repens* and separated lobes of the clover *Oxalis atropurpurea* were used for microscopy. The leaves were excised from the plants and mounted directly on slides with the abaxial surface facing upwards, covered with fresh water and covered with a coverslip. The microscopes used were the Zeiss Axio-Lab A1 and the Zeiss discovery V20 stereo microscope coupled with the Axiocam ERc5s and Icc1 cameras under the management of the Zeiss Axiovision software. The magnifications used varied among the plants. Thus, for Tradescantia pallida, Tradescantia zebrina, Tradescantia minima and Callisia repens, the magnifications 15 × were used in the stereomicroscope and 50×, 100×, 200× and 400 × in the Axio-Lab A1 microscope. For the plants *Ctenante oppenheimiana, Oxalis atropurpurea*, the magnifications of 50 ×, 100 ×, 200 × and 400 × obtained with the objectives of the AxioLab A1 microscope were used.

### 1.2 Measure of distances between stomata

The microscopic images were processed using Matlab image processing toolbox. The stomates were identified manually in the images, their centroids (*x, y*) were identified through an algorithm which found the central coordinates of each one of the manually marked dots. The distance between each pair of points in the image is given by the following equation, where d is the distance between two centroids (i,j) of the stomata.

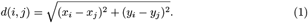

Images with magnifications of 100 × were used. A distance matrix between each pair of centroid stomata at image is calculated as illustrated in Fig. 3.

## 2. Results

### 2.1 Plant microscopy

Samples of microscopic images of six plant species with purple abaxial leaf are shown in the following figure. The optical microscopy images obtained from *Tradescantia pallida, Tradescantia zebrina, Callisia repens, Tradescantia minima, Ctenanthe oppenheimiana* and *Oxalis atropurpurea* show green stomata in extant leaves.

The microscopic magnification values used to collect the images of the Figure 2, give the dimension of the size of the stomata in the studied species. For example, the length of the stomata from *Tradescantia pallida*, the largest among all the plants studied, is on the order of more than 10× the stomata of the plant *Oxalis atropurpurea*, the smallest. Although all plants have the pavement color contrasting with the color of the stomata, the color of the abaxial face has plastic behaviour, especially in the specie *Oxalis atropurpurea* and *Tradescantia minima* in which they varied from green to purple according to environmental variations [1].

### 2.2 Morphometrics

Depending on the species and the environmental conditions stomata range in size from about 10 to 80*µ*m in length and occur at densities between 5 and 1,000 *mm*^*−*2^ of epidermis [15].

In spite of this wide variability there is a general relationship between density and size for different plant groups (grasses and non-grasses)and fossil leaves. [8].. Simulations with a stomata model of gaseous exchange indicate that the average relationship has virtually the same trend to one in which changes in density and size are exactly compensatory, in terms of *CO*_2_ and water vapour exchange [2, 8].

The distance measurements of interstomatic distances according illustrated at figure 1 quantify how the plasticity of the leaf can be measured from the distribution of these cells in the epidermis. The measurements of the distances between neighboring stomata × stomatal densities can be evidenced in the graphs of Figure 3, which show the dispersion of the density values as a function of the mean distance between the stomata neighbors for the six species of purple plants.

**Figure 1.**
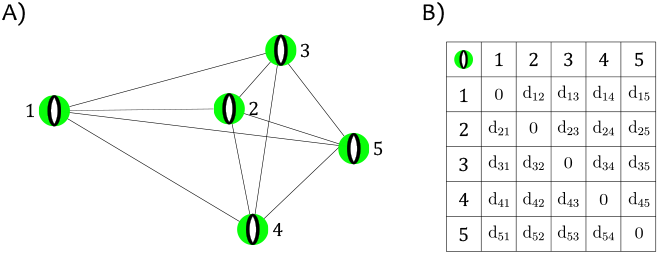
A) Illustration of five stomates (green) connected with each other by lines. B) Distance matrix of each stomata pair. The matrix index represent each stomata, and *d*(*i, j*) is the distance between each pair of stomata found in A).

**Figure 2.**
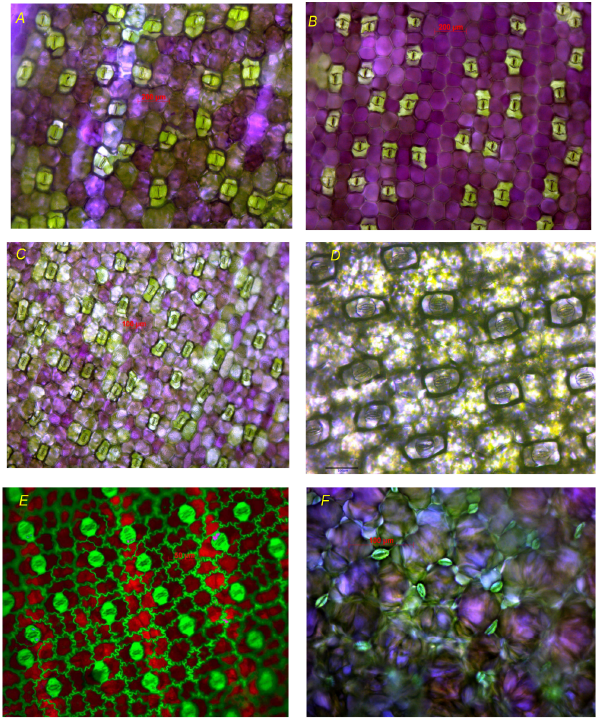
Optical microscopy images shown green stomata distributed over the abaxial surface of leaves. A) Image from *Tradescantia pallida* magnified 50× B) Image from *Tradescantia zebrina* magnified 50× C) Image from *Callisia repens* magnified 50× D) Image from *Tradescantia minima* magnified 100×. E) Image from *Ctenanthe oppenheimiana*, magnified 200× F) Image from *Oxalis atropurpurea* magnified 200×.

**Figure 3.**
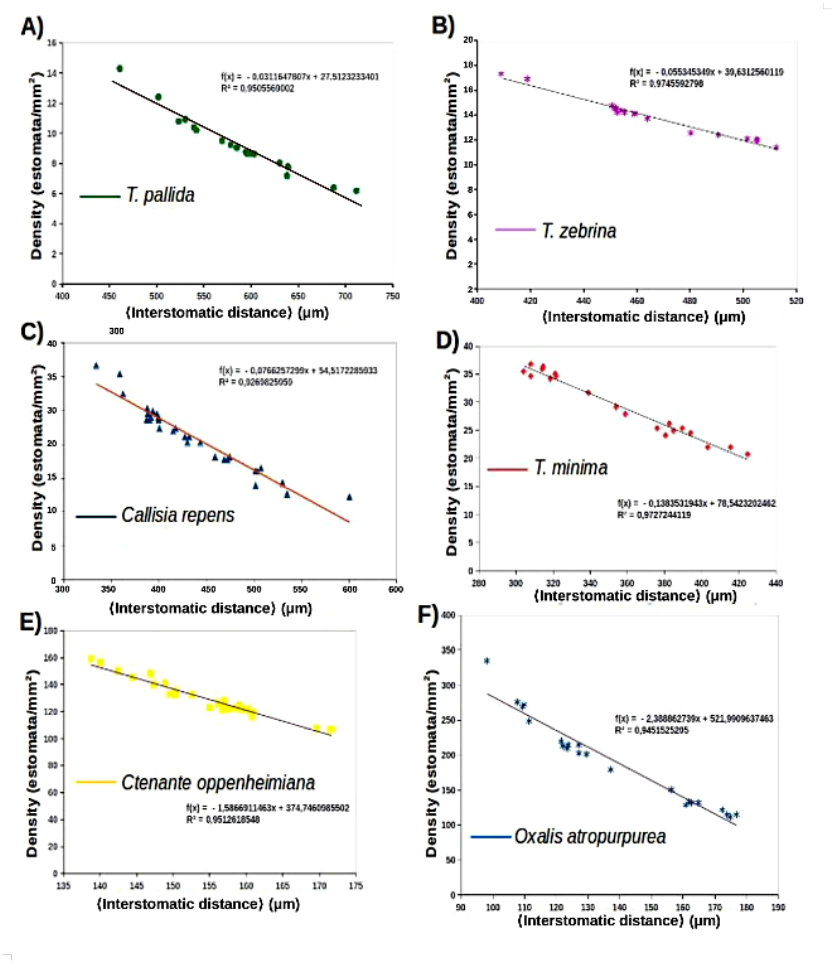
Dispersion of the mean distance values between neighboring stomata as a function of the mean stomatal density for six plants, plots from top to bottom: A) *T. pallida* B) *Tradescantia zebrina* C) *Callisia repens*. D) *T. minima* E) Ctenanthe oppenheimiana. F) *Oxalis atropurpurea*.

The results indicate that the distances between neighboring stomata are inversely proportional to their stomatic densities and vary linearly in each species independently. These measurements are, therefore, direct evidence of plasticity as regards the variation of stomata distribution on leaf epidermis. When transposing the mean values of the measurements indicated in the graphs shown above, for a single graph that quantifies the relation mean distance × mean density to 6 plant species studied here, it can be observed the similar log-normal trend line as observed to relation stomatic length in function of stomatic density showed in previous works [2, 8] (see fig 4).

**Figure 4.**
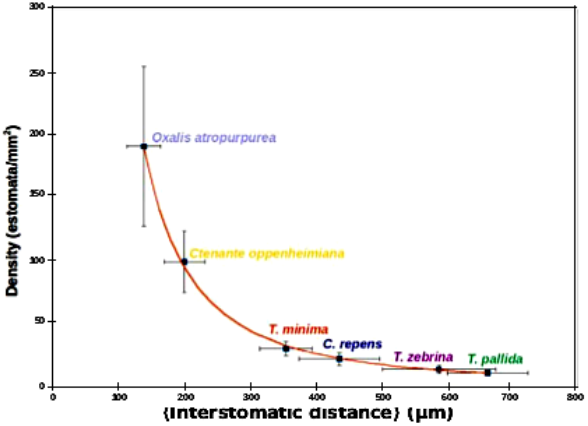
Dispersion of the mean interstomatic distance × mean stomatic density for *Tradescantia pallida, Tradescantia zebrina, Callisia repens, Tradescantia minima, Ctenanthe oppenheimiana* and *Oxalis atropurpurea*. The solid curve is log-normal, y = 2260026.3 × x^*−*1.90^, *r*^2^ = 0.99 The error bars indicate the standard deviation of the measurements in relation to the mean values of distances between stomata and stomatic densities by species. (N = 5 plants per species, 3 images per plant)

The measures of distance between stomata are used as a parameter to measure phenotypic plasticity. The results shown here indicate that this measure quantifies the dispersion of the stomata and precisely measures its distribution on the leaf. Plants with a purple abaxial surface exhibit a natural color contrast with stomata, which are usually green due to the large amount of chlorophyll and chloroplasts present within the guard cell. Therefore the measurement of distance between stomata in purple abaxial surface plants, seems to be a powerful strategy for observation of changes in the physiology of plants, generated by changes in the environment. Thus ecophysiological aspects in plants will be approached more accurately from the estimation in distribution of stomata on the leaves. It is known that the stomata feel the environmental changes, being natural sensors of the plant the conditions of the environment, at the same time that they are responsible for the climate change [8]. Thus the strategy of stomata morphometry in purple plants, showed here, may provide valuable answers to the understanding of how plants respond to climate change.

## Discussion

The stomata are modified cells responsible to plant gas exchange, consequently play fundamental roles in photosynthesis and carbon assimilation by plants. Consequently, they are fundamental in the flows of carbon and water on planet Earth [4, 8, 12]. Therefore, quantitative information on the morphology of stomata in the leaf with data on relationships between stomatic density, stomata size and the environmental plasticity of these characters are important probes to understand how plants perceive and feel environmental changes. We observed in this work that microscopic images of purple abaxial epidermis of living plants show stomata that are completely visible and discernible from the rest of the epidermis due to the contrast between the purple colors of the epidermis and the green stomata (see figure 2). The contrast green/purple is very striking in the plant of the species *Ctenanthe oppenheimiana* (see figure 2 E)). It allowed us to count the distribution of these cells, determine their size and measure the distance between them in the microscopic images of 6 plants with purple abaxial leaves. The measurements revealed similar morphological relationships to those already observed in previous works [2, 8], which identify an inverse log-normal relationship between stomata density and stomata size as a general law applicable to a large number of plants including fossil plants. These studies also show that stomatic density is highly variable when plants grow in different concentrations of *CO*_2_ and that this characteristic is striking in plants grown in different geological eras on Earth, especially during the Phanerozoic, when the concentrations of *CO*_2_ in the atmosphere are different [2]. The measurements in living plants with purple abaxial epidermis made here show a correlation coefficient with a log-normal curve much higher than that observed in previous studies (*R*^2^ > 0.9) (See figure 4). This probably occurs because few plants were used here, with stomatal density values with larger intervals between them. Furthermore the measurements were made in living plants and without histological processing of the leaf. These data confirm the simulations with a stomatal model of gas exchange made in previous studies [2] and confirm the inverse relationship between stomata density and stomata size as a log-normal relationship.

The compilation of the distance measurements between stomata through a quadratic matrix (see figure 1) proved to be a powerful tool for measuring the plasticity of stomatal distribution in the leaf. This is confirmed by the inverse log-normal relationship found between the mean distance between stomata and stomatic density among different plant species (see figure 4). It is interesting to note that this relationship is linear for measurements made in a single species and logarithmic when observed between different species (see figures 3 and 4). These results show that the strategy of using plants with purple abaxial epidermis as sensors of environmental change is achievable through stomata distribution measurements using interstomatic distances.

